# Land-surface evapotranspiration derived from a first-principles primary production model

**DOI:** 10.1101/2021.06.23.449361

**Authors:** Shen Tan, Han Wang, Iain Colin Prentice, Kun Yang

## Abstract

We propose an application of eco-evolutionary optimality theory in the context of monitoring and modelling physical land-surface processes. Evapotranspiration (ET) links the water and carbon cycles in the atmosphere, hydrosphere, and biosphere. We develop an ET modelling framework based on the hypothesis that canopy conductance acclimates to plant growth conditions so that the total costs of maintaining carboxylation and transpiration capacities are minimized. This is combined with the principle of co-ordination between the light- and Rubisco-limited rates of photosynthesis to predict gross primary production (GPP). Transpiration (T) is predicted from GPP via canopy conductance. No plant type- or biome-specific parameters are required. ET is estimated from T by calibrating a site-specific (but time-invariant) ratio of modelled average T to observed average ET. Predicted seasonal cycles of GPP were well supported by (weekly) GPP data at 20 widely distributed eddy-covariance flux sites (228 site-years), with correlation coefficients (*r*) = 0.81 and root-mean-square error (RMSE) = 18.7 gC/week and Nash-Sutcliffe efficiency coefficient (NSE) = 0.61. Seasonal cycles of ET were also well supported, with *r* = 0.85, RMSE = 5.5 mm week^−1^ and NSE = 0.66. Estimated T/ET ratios (0.52–0.92) showed significant positive relationships to radiation, precipitation and green vegetation cover and negative relationships to temperature and modelled T (*r* = 0.84). Although there are still uncertainties to be improved in the current framework, particularly in estimating T/ET, we see the application of eco-evolutionary principles as a promising direction for water resources research.

**Highlights:**

- Building an evapotranspiration estimation framework based on *a priori* primary productivity model (the P model).
- Assessing the contribution of environmental indicators to the ratio of transpiration to evapotranspiration.
- Proving the reliability of this approach to estimate evapotranspiration.

## 1. Introduction

Evapotranspiration (ET) is a key process in the global terrestrial water cycle (Bai and Liu, 2018). ET comprises the biotic process of transpiration (T) via stomata, and the abiotic processes of evaporation from wet leaves (interception) and bare soil. ET is a core object for regional water management (Bastiaanssen et al., 2000; Zeng et al., 2019) and the mitigation of excessive water consumption (Fisher et al., 2017). It mediates energy exchange at the land-atmosphere interface and thereby influences regional and global climate (Trenberth et al., 2009). Transpiration can indicate the growth status of both natural and cultivated vegetation canopies due to the close coupling between carbon uptake and water transpired (Ershadi et al., 2014). Accurate modelling and monitoring of ET are thus important for multiple applications in ecology, hydrology and climate science.

Many ET models depend on remote sensing inputs (Mu et al., 2007, Wu et al., 2012). Apart from empirical relationships employed in early studies (Moran et al., 1994, Nagler et al., 2013) and more recent machine-learning approaches (Jung et al., 2009, Torres et al., 2011, Elnashar et al., 2021), ET is generally predicted based on the land-surface energy balance. A conventional approach is to calculate the latent heat flux representing the energy flux during the vaporization of water as the residual of the surface energy balance equation (Bastiaanssen et al., 1998; Su, 2002; Norman et al., 1995). Sensible heat flux is commonly predicted based on its relationship with the temperature gradient between the atmosphere at reference height and the land surface, which can be simulated using remotely observed land surface temperature (LST) (Lagouarde et al., 1991). However, remotely sensed signals suffer from extensive cloud contamination. Due to the extreme sensitivity of sensible heat flux to LST, and LST to instantaneous climate conditions, the robustness of monitoring time-series ET by this strategy may be compromised by input data quality (Tan et al., 2019).

The Penman-Monteith (PM) equation provides an alternative, more direct approach to estimate the latent heat flux and ET. This equation has also been applied widely to calculate ET (Granger et al., 1989; Carlson et al., 1991; Allen et al., 1998; Cleugh et al., 2007; Mu et al., 2007; Leuning et al., 2008). Its application depends on specifying surface conductance. Jarvis (1976) suggested that surface conductance could be represented for different land-cover types by combining its empirical responses to meteorological variables including air temperature, downward shortwave radiation, and soil moisture. This approach has been applied globally (Mu et al., 2007, Zhang et al., 2019). However, ‘Jarvis-type’ modelling of surface conductance entails substantial uncertainties because of the need to calibrate multiple parameters for different land cover types.

Modern approaches to estimating surface conductance and ET in Earth System models (ESMs) rely on the intrinsic coupling between photosynthesis and transpiration, both regulated by stomata (Wong et al., 1979). Ball and Berry (1987) proposed an empirical model of stomatal conductance as a function of photosynthesis and relative humidity (RH). Leuning (1995) developed this idea further, noting that vapour pressure deficit (VPD) rather than RH is the driving force of ET (Leuning et al., 2008). These models have been applied widely at a global scale (Kowalczyk et al., 2006, Oleson et al., 2010; Jiang et al., 2016; He et al., 2018; Zhang et al., 2019). Medlyn et al. (2011) introduced a new stomatal conductance model based on the Cowan and Farquhar (1977) hypothesis on optimal stomatal behavior. This new model has features in common with the empirical models of Ball and Berry (1987) and Leuning (1995), and provides a partial theoretical basis for them. However, all these models require the calibration of parameters that vary substantially among biomes and regions (Lin et al., 2015; Knauer et al., 2018) and an assumption of their constancy under changing environmental conditions. The multiplicity of poorly known parameters, combined with the questionable assumption that they are constant, is likely to limit the usefulness of current ET models for climate-change applications (Yang et al., 2019). Moreover, current ESMs show a pervasive bias, systematically underestimating the importance of transpiration – to a greater or lesser extent in different models – due to systematic errors in model representations of light absorption in plant canopies (Lian et al., 2018).

It would be highly desirable, therefore, to develop universal models of photosynthesis and stomatal conductance, applicable across biomes without re-calibration. A recently developed model for gross primary production (GPP) (the P model: Wang et al., 2017, Stocker et al., 2020) provides such a way to avoid the limitations of current models. At the core of the P model is a description of optimal stomatal behaviour as a function of environment that is equally applicable to all C_3_ plants (Prentice et al., 2014). The P model has been used to diagnose the response of the terrestrial carbon cycle to changes in climate and atmospheric CO_2_ (e.g. Keenan et al. 2016; Stocker et al., 2020); as part of a global monitoring system for GPP (https://terra-p.vito.be); and as the basis for a generic scheme to predict wheat yields (Qiao et al., 2020). Pérez◻Priego et al. (2018) combined the P model and an empirical canopy conductance model to partition observed ET into transpiration and evaporation, suggesting the potential to employ the P model globally to estimate the transpiration component of ET.

Here we propose a modelling framework for ET based on the PM equation and the P model’s estimation of canopy conductance. This framework combines optimality theory with remote sensing data. We use site-level observations to evaluate the model.

## 2 Materials and methods

### 2.1 A general framework for modelling ET

The central operation in our proposed ET modelling framework is the estimation of canopy conductance, based on GPP and the ratio of leaf-internal to ambient CO_2_ partial pressure (*c*_i_/*c*_a_, denoted by *χ*), both of which are calculated by P model. This calculation is independent of plant functional types, except for the necessary distinction between C_3_ and C_4_ plants. The predicted GPP and *χ* values are used to estimate canopy conductance to CO_2_, which is multiplied by 1.6 to yield the estimated average canopy conductance to water vapour (*G*_s_, mol m^−2^ s^−1^). We then estimate T with the PM equation driven by the predicted canopy conductance and meteorological variables. A weekly time step is adopted for the prediction of GPP, T and ET because the theory behind the P model is based on leaf-level acclimation of photosynthetic parameters to environment, which occurs over a time scale of weeks.

The ratio T/ET is constrained by local environment due to water limitation under dry conditions and energy limitation under wet conditions. There is evidence that this ratio is temporally conservative without disturbance on the local environment (Paschalis et al. 2018), perhaps because local wetness condition simultaneously controls transpiration and soil evaporation. Here, accordingly, we calculate site-specific (but time-invariant) values of T/ET from the predicted T and observed ET, allowing us both to test our simulations of the seasonal cycle of ET and to explore the potential environmental dependencies of this ratio (Fig. 1).

**Fig. 1.**
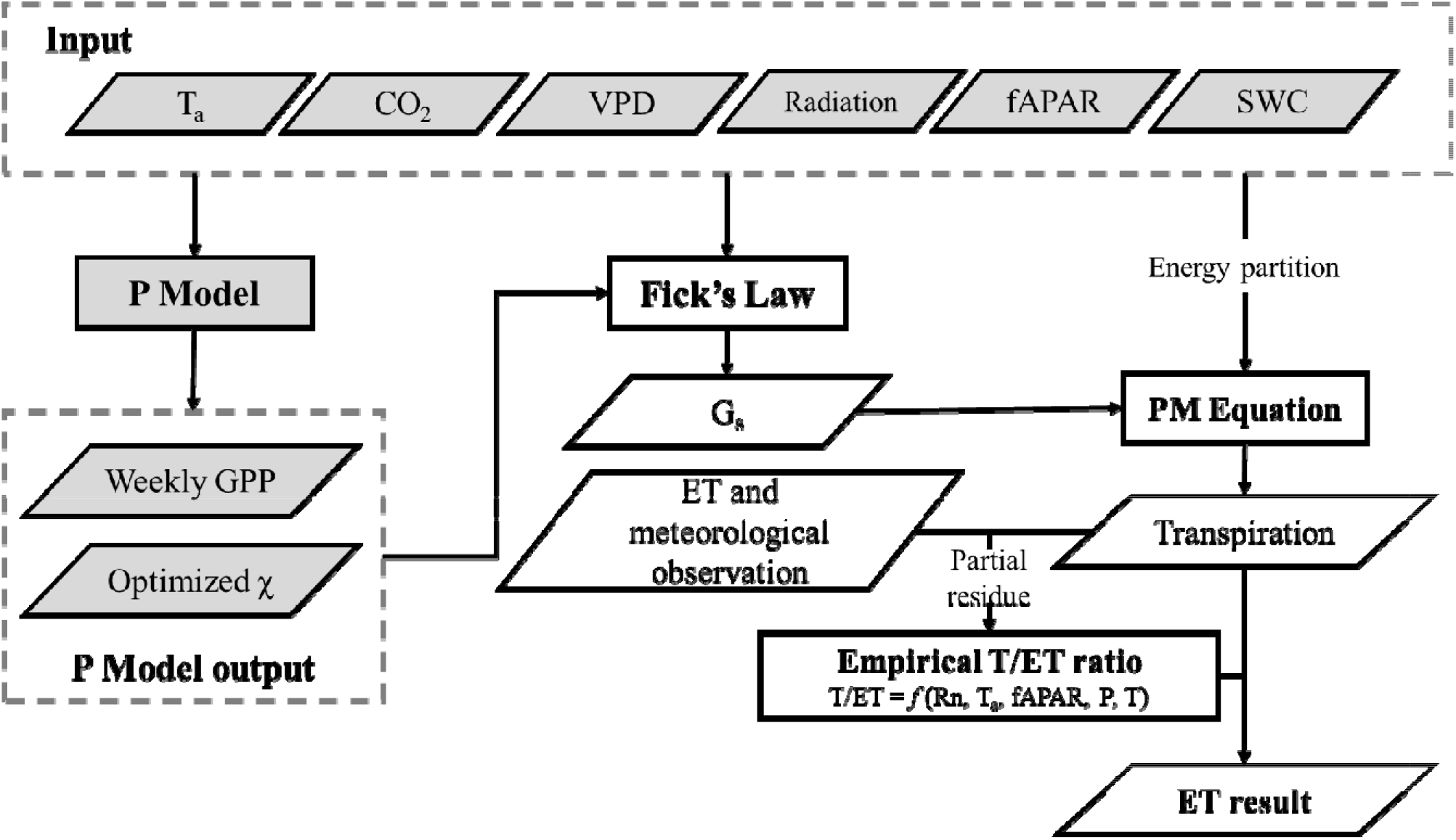
Flowchart of the P model based ET prediction. Items in grey have been validated by previous P model studies, summarized in Section 2.1.1. “Radiation” in the input block includes the four components of net radiation (Rn), and PPFD (photosynthetic photon flux density). SWC is the volumetric soil water content, which is an input to the P model. Canopy conductance is calculated by Fick’s law, which requires simulated weekly GPP and *χ* (ratio of *c*_i_/*c*_a_) as input. The ratio T/ET is required for calculating ET from modelled transpiration, and is fitted by ET and meteorological observation. In the empirical function, *T*_a_ is air temperature in °C, fAPAR is the fraction of absorbed photosynthetically active radiation, and P is annual precipitation in mm.

#### 2.1.1 Predicting GPP and χ using the P model

The P model is an extension of the FvCB biochemical model of C_3_ photosynthesis (Farquhar et al., 1980) to account for optimal acclimation of photosynthetic capacities and stomatal behaviour (Wang et al., 2017, Stocker et al., 2020: the reader is referred to these publications for the detailed equations and their derivation). The instantaneous rate of photosynthesis according to the FvCB model is the lesser of the electron transport-limited rate (*A*_J_) and the carboxylation-limited rate (*A*_C_). Both rates are limited by *c*_i_, and therefore depend on the product of *c*_a_ and *χ*. Based on the concept of eco-evolutionary optimality (Franklin et al., 2020), the *least-cost hypothesis* states that plants minimize the total costs of maintaining carboxylation capacity and transpiration through the regulation of stomatal conductance (Wright et al., 2003; Prentice et al., 2014). This hypothesis leads to a prediction of optimal *χ* as a function of VPD, temperature and elevation (Prentice et al., 2014; Wang et al., 2017).

A second optimality hypothesis, the *coordination hypothesis*, states that on a weekly to monthly time scale, *A*_J_ and *A*_C_ converge (Chen et al. 1993; Haxeltine and Prentice 1996; Maire et al., 2012) by acclimation of the maximum rate of carboxylation (*V*_cmax_) (Smith et al., 2019). The maximum rate of electron transport (*J*_max_) is observed to vary in parallel to *V*_*c*max_. Optimal *J*_max_ is assumed to maximize the difference between the benefit (*A*_J_) and cost of maintaining a certain level of *J*_max_ (Wang et al., 2017; Smith et al., 2019). Wang et al. (2017) showed that the least-cost and coordination hypotheses, together with this principle for the optimization of *J*_max_, lead to a closed-form expression for GPP. Despite its basis in the FvCB model of instantaneous photosynthesis (which implies a non-linear, saturating response to light), this model has the mathematical form of a light use efficiency (LUE) model: in other words, accumulated GPP for C3 plant over the acclimation time scale is proportional to accumulated absorbed photosynthetic photon flux density (PPFD). *V*_cmax_ and *J*_max_ do not have to be specified, because their optimal values are implicitly calculated by the model.

We modified the P model to predict C_4_ as well as C_3_ photosynthesis based on the theory by Collatz et al., (1992). Given that phosphoenolpyruvate carboxylase (the initial carboxylating enzyme in the C_4_ pathway) has a higher affinity for CO_2_ than Rubisco (the primary carboxylating enzyme of the C_3_ pathway), we assumed that C_4_ photosynthesis is not limited by the intercellular CO_2_ concentration (Sayre et al., 1979). The model as applied here also includes the generic soil-moisture limitation function described by Stocker et al. (2020) and temperature dependencies of the intrinsic quantum efficiency of photosynthesis, based on measurements by Bernacchi et al. (2003) for C_3_ plants (Stoker et al., 2020) and Kubien et al. (2003) for C_4_ plants (Cai & Prentice, 2020). The soil moisture function requires an estimate of a climatological aridity index, which was calculated for each site using in-situ observation and the Priestley-Taylor equation (Priestley and Taylor, 1972). The full model code is available at https://github.com/stineb/rpmodel.

#### 2.1.2 Canopy conductance

The diffusion of both water vapour and CO_2_ through stomata can be described by Fick’s law. CO_2_ diffusion into the leaf is driven by the difference between the ambient and leaf-internal concentrations. We scale up from leaf to canopy level in the simplest possible way via the “big-leaf” approximation. Thus, canopy stomatal conductance to water vapour is given by:

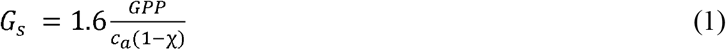

where the factor 1.6 is the ratio of the molecular diffusivities of water and CO_2_. For C_4_ plants, we set *χ* in equation (1) to a typical value of 0.45 (Farquhar et al., 1989).

#### 2.1.3 Transpiration

Total latent heat flux (*λE_t_*) can then be decomposed into contributions from transpiration (*λE_c_*) and evaporation (*λE*):

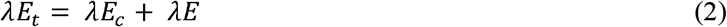

where *λ* is the latent heat of evaporation of water (MJ kg^−1^). *λE*_*c*_ was calculated using the Penman-Monteith equation:

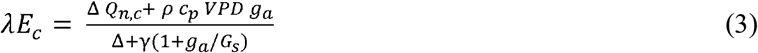

where Δ is the slope of the curve relating saturation water vapour pressure to air temperature (kPa K^−1^) and *Q_n,c_* is the available energy (net radiation) intercepted by the canopy (W m^−2^), i.e., a fraction of the total net radiation minus soil heat flux (*R*_n_ – *G*). Because shortwave radiation is generally the largest part of net radiation, this fraction was estimated by Beer’s Law as *Q_n,c_*/*Q_n_* = 1 – exp (−*k LAI*), where *k* is the extinction coefficient, assumed constant at 0.5. This has a similar mathematical form to the calculation of fAPAR from LAI (leaf area index) (Gan et al., 2018; Zhang et al., 2019). *ρ* in equation (3) is the density of air (kg m^−3^), *c_p_* is the heat capacity of dry air (J kg^−1^ K^−1^), *G_s_* is calculated by equation (1), and *g_a_* is the aerodynamic conductance calculated by the simple model of Thom (1972):

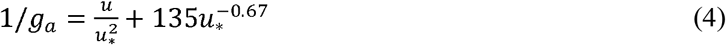

where *u* is the wind speed (m s^−1^) and *u*_*_ (m s^−1^) is the friction velocity, which is obtained from flux observations. The even simpler equation recommended by FAO (Allen et al., 1998) was used when no information on *u*_*_ was available:

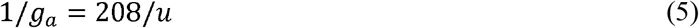

#### 2.1.4 Site-specific ratio of transpiration to total evapotranspiration

Fatichi et al. (2017) used a detailed, mechanistic ecosystem model that simultaneously solves for water, energy, and carbon exchanges over the land surface to analyse global variation in simulated T/ET. Their results suggested that T/ET is globally constrained to 70 ± 9% with only modest effects of climate or vegetation type – although under some extreme conditions, such as drip-irrigated farmland, the ratio could lie outside this range (Kool et al., 2014). Here we assumed a constant T/ET ratio at each site, and estimated this ratio using a priori relationship between modelled transpiration as described above (equations 1–5) and the observed ET and meteorological conditions.

We then estimated partial effects (effects of each predictor variable with the others held constant at their median values) in a multiple regression to analyse how T/ET ratio varies with environment. Five indicators (*R*_n_, fAPAR, *T*_a_, *P*, and modelled transpiration) were selected as potential predictors. The first four of these indicators are the inputs to the empirical soil evaporation model of Zhang et al. (2010). Transpiration was selected to describe the contribution of environmental water supply, since soil water content information is not available at all flux sites.

### 2.2 Sites and in situ observations

Twenty sites with 228 observation-years of flux data (including sensible heat flux, latent heat flux and soil heat flux) from the FLUXNET2015 Tier 1 data (https://fluxnet.fluxdata.org) were used. We employed a site selection criterion that the energy balance ratio (Rn – G)/(+ H) must lie between 0.8 to 1.2 (where H represents sensible heat flux). These sites span eight main land cover types (Table 1, Fig. 2). Weekly measurements of *T*_a_, wind speed, the four components of radiation (downward and upward long-wave and short-wave radiation), soil water content and ambient CO_2_ concentration are recorded at all these sites, and were used to drive the P model. Daily ecosystem carbon exchange was partitioned into GPP and ecosystem respiration using the half-hourly night-time separation method of Reichstein et al. (2005). Unreliable records were filtered out before weekly averaging by imposing the following requirements: (1) available energy (net radiation minus soil heat flux) > 0, which is a pre-requisite for evapotranspiration; (2) average air temperature > 5 ◻ and GPP > 0; (3) precipitation = 0; (4) LAI > 0; (5) less than 10% fAPAR is ineffective in the focused year (e.g. GF-Guy in 2014 was heavily contaminated by cloud).

**Table 1.** Information on the selected flux sites. **Lon** = longitude (°), **Lat** = latitude (°), **Ele** = elevation (m). **MAT** = mean annual temperature (C), **MAP** = mean annual precipitation (mm). Biome type codes are the same as in Fig. 2.

| **D** | **Site Name** | **L on** | **L at** | **E le** | **I GBP** | **Data Range** | **M AT** | **M AP** |
| --- | --- | --- | --- | --- | --- | --- | --- | --- |
|  | AU-D | 13 | -1 | 4 | S | 2008-2 | 2 | 9 |
|  | as | 1.39 | 4.16 | 70 | AV | 014 | 7.2 | 76 |
|  | AU-T | 14 | -3 | 1 | E | 2001-2 | 1 | 1 |
|  | um | 8.15 | 5.66 | 200 | BF | 014 | 0.7 | 159 |
|  | BR-S | -5 | -3 | 1 | E | 2000-2 | 2 | 2 |
|  | a3 | 4.97 | .02 | 00 | BF | 004 | 6.1 | 043 |
|  | CA-M | -9 | 5 | 2 | E | 1994-2 | - | 5 |
|  | an | 8.48 | 5.88 | 59 | NF | 008 | 3.2 | 20 |
|  | CA-T | -8 | 4 | 1 | E | 2009-2 |  | 1 |
|  | p3 | 0.35 | 2.71 | 84 | NF | 014 | 8 | 036 |
|  | CN-C | 12 | 4 | 7 | M | 2003-2 | 2 | 6 |
|  | ha | 8.1 | 2.4 | 66 | F | 005 | .2 | 64 |
|  | CN-D | 11 | 2 | 3 | E | 2003-2 | 1 | 1 |
|  | in | 2.54 | 3.17 | 08 | BF | 005 | 9.6 | 618 |
|  | CN-Q | 11 | 2 | 1 | E | 2003-2 | 1 | 1 |
|  | ia | 5.06 | 6.74 | 09 | NF | 005 | 9 | 467 |
|  | DE-T | 13 | 5 | 3 | E | 1996-2 | 8 | 8 |
|  | ha | .57 | 0.96 | 85 | NF | 014 | .2 | 43 |
|  | DK-S | 11 | 5 | 4 | D | 1996-2 | 8 | 6 |
| 0 | or | .64 | 5.49 | 0 | BF | 014 | .2 | 60 |
|  | FI-Hy | 24 | 6 | 1 | E | 1996-2 | 3 | 7 |
| 1 | y | .29 | 1.85 | 81 | NF | 014 | .8 | 09 |
|  | FR-Gr | 1. | 4 | 1 | C | 2004-2 | 1 | 6 |
| 2 | i | 95 | 8.84 | 25 | RO | 014 | 2 | 50 |
|  | GF-G | -5 | 5. | 4 | E | 2004-2 | 2 | 3 |
| 3 | uy | 2.92 | 28 | 8 | BF | 014 | 5.7 | 041 |
|  | IT-Ro | 11 | 4 | 1 | D | 2002-2 | 1 | 8 |
| 4 | 2 | .92 | 2.39 | 60 | BF | 012 | 5.2 | 76 |
|  | RU-S | 12 | 6 | 2 | D | 2012-2 | - | 2 |
| 5 | kp | 9.17 | 2.26 | 46 | NF | 014 | 1.1 | 90 |
|  | US-A | -9 | 3 | 3 | C | 2003-2 | 1 | 8 |
| 6 | rm | 7.49 | 6.61 | 14 | RO | 012 | 4.8 | 43 |
|  | US-H | -7 | 4 | 3 | D | 1991-2 | 6 | 1 |
| 7 | a1 | 2.17 | 2.54 | 40 | BF | 012 | .6 | 071 |
|  | US-N | -9 | 4 | 3 | M | 2001-2 | 1 | 7 |
| 8 | e1 | 6.48 | 1.17 | 61 | AI | 013 | 0.1 | 90 |
|  | US-P | -9 | 4 | 4 | M | 1995-2 | 4 | 8 |
| 9 | Fa | 0.27 | 5.95 | 72 | F | 014 | .3 | 23 |
|  | ZA-K | 31 | -2 | 3 | S | 2000-2 | 2 | 5 |
| 0 | ru | .5 | 5.02 | 59 | AV | 013 | 1.9 | 47 |

**Fig. 2.**
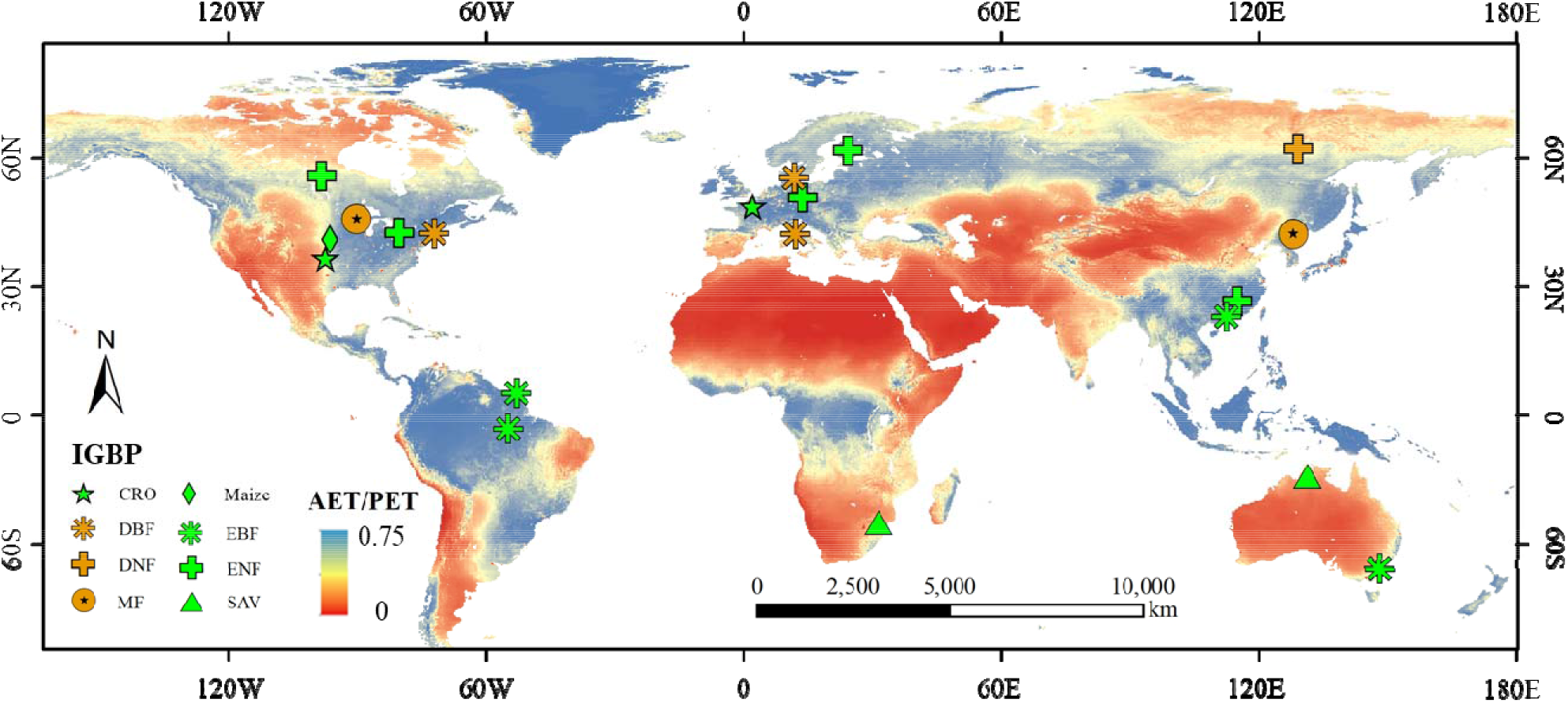
Map of flux sites. Twenty sites from eight major biomes, according to the International Geosphere-Biosphere Programme (IGBP) classification. **CRO** = crop, **DBF** = deciduous broadleaf forests, **DNF** = deciduous needleleaf forests, **MF** = mixed forests, **EBF** = evergreen broadleaf forests, **ENF** = evergreen needleleaf forests, **SAV** = savanna, **MAI** = maize. The map background is averaged ratio of mean ratio of actual evapotranspiration (AET) dividing potential evapotranspiration (PET) through the year of 1990 to 2015 from GLDAS (Global Land Data Assimilation System, v2.1) (Rodell et al., 2004).

### 2.3 Remote sensing observations

Remotely sensed fAPAR is required as an input by the P model. Since some of the flux records started in the early 1990s, we used the fAPAR product from NOAA CDR AVHRR with a 0.05° spatial resolution in order to maintain consistency of the model input (Claverie and Vermote, 2014). fAPAR time series were smoothed by a Savitzky–Golay filter to eliminate high-frequency noise (Chen et al., 2004).

## 3 Results

### 3.1 Predicting weekly GPP using the P model

Weekly predicted GPP was well supported by observations (Fig. 3 and Table 2). The correlation coefficient (*r*) based on all records was 0.81, the RMSE (root mean squared error) is 18.7 gC/week, and the NSE (Nash-Sutcliffe efficiency coefficient) is 0.61. At site scale, *r* ranged from 0.48 (at BR-Sa3) to 0.92 (at CN-Cha), and RMSE from 6.2 gC/week (at RU-Skp) to 24.8 gC/week (at US-Ne1). This good performance is consistent with previous P model evaluation studies (Wang et al., 2017; Stocker et al., 2018, 2020; Qiao et al., 2020). The good performance at the maize site (US-Ne1) supports its extension to C_4_ vegetation (RMSE = 24.8 gC/week, and a 24.5% of relative root mean squared error, denoted as RRMSE). The model tends to slightly underestimate GPP (fitting slope is 0.81); this bias can be removed by calibrating a single, global parameter representing the efficacy with which absorbed light is used in photosynthesis (Stocker et al., 2020), but we did not apply any such *a posteriori* adjustment. The underestimation is probably due at least in part to mismatches of the flux tower footprints with the AVHRR fAPAR grids (Kljun et al., 2015). Heterogeneity inside the target RS pixel (0.05° × 0.05°), but not in the tower footprint, especially patches of bare soil, would be expected to reduce the measured fAPAR by increasing reflectance in the red band, leading to an underestimation of absorbed radiation.

**Table 2.** Statistical information on the GPP evaluations on a weekly timestep. Slope: coefficients of linear fits, *r*: correlations between observed and predicted values, RMSE: root mean squared error of prediction, RRMSE: relative root mean squared error.

| Site Name | GPP estimation result |  |  |
| --- | --- | --- | --- |
| | $r$ | RMSE (g C/week) | RRMSE (%) |
| AU-Das | 0.75 | 7.4 | 23.5 |
| AU-Tum | 0.59 | 16.1 | 26.6 |
| BR-Sa3 | 0.48 | 15.8 | 25.0 |
| CA-Man | 0.77 | 10.3 | 34.7 |
| CA-TP3 | 0.61 | 20.3 | 44.1 |
| CN-Cha | 0.92 | 9.2 | 16.4 |
| CN-Din | 0.78 | 7.8 | 25.0 |
| CN-Qia | 0.87 | 8.2 | 21.6 |
| DE-Tha | 0.75 | 14.6 | 28.4 |
| DK-Sor | 0.89 | 16.1 | 24.2 |
| FI-Hyy | 0.89 | 11.7 | 27.4 |
| FR-Gri | 0.61 | 25.3 | 34.4 |
| GF-Guy | 0.49 | 24.1 | 33.3 |
| IT-Ro2 | 0.63 | 16.9 | 34.9 |
| RU-Skp | 0.82 | 6.2 | 24.6 |
| US-Arm | 0.68 | 7.7 | 22.8 |
| US-Ha1 | 0.84 | 19.3 | 33.0 |
| US-Ne1 | 0.86 | 24.8 | 24.5 |
| US-PFa | 0.88 | 8.5 | 20.7 |
| ZA-Kru | 0.65 | 9.3 | 24.2 |

**Fig. 3.**
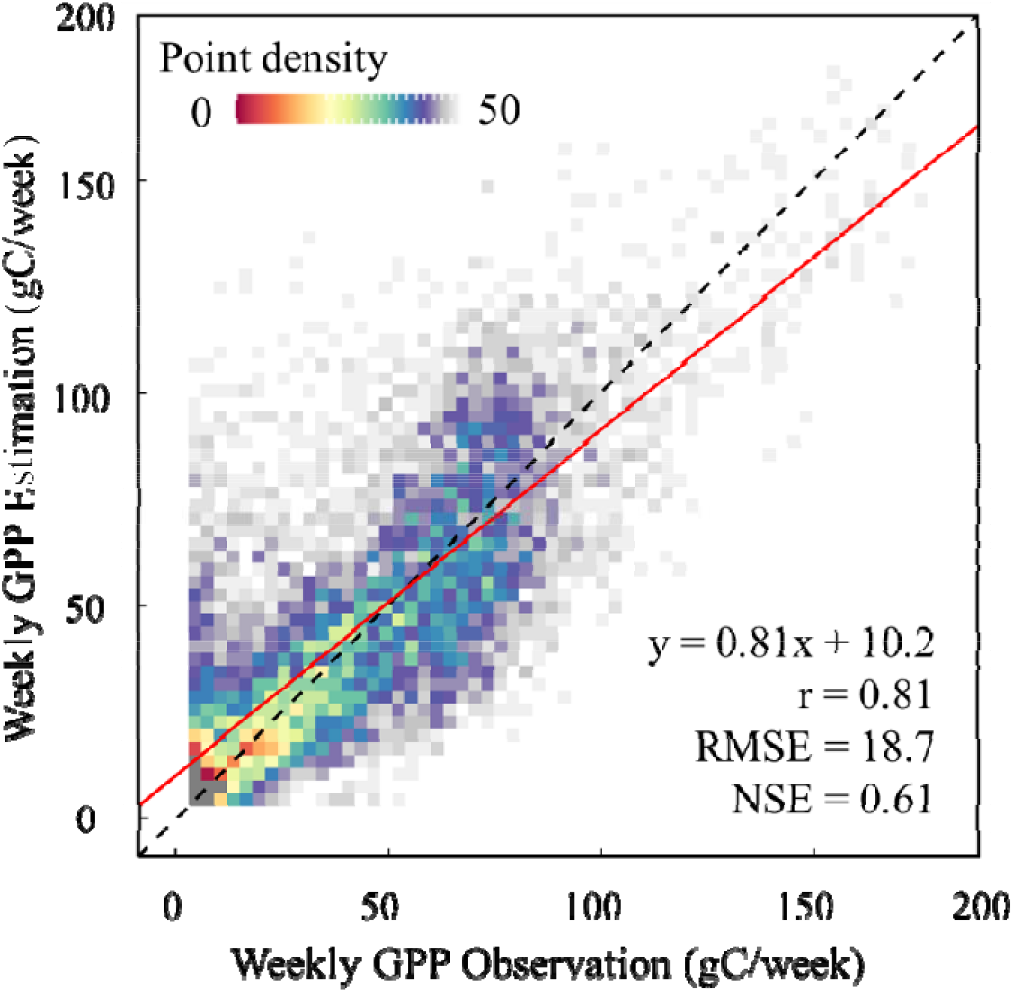
Comparison of estimated GPP against observations at a weekly time scale. The red line is the linear fit; the dashed line is the 1:1 line. Colours represents the density of points.

### 3.2 Estimating T/ET ratio

The fitted slope in Table 3 indicates that the ratio T/ET ranged from 0.52 (FR-Gri) to 0.92 (CA-Man). The five selected predictors of this ratio (net radiation, fAPAR, precipitation, air temperature and modelled transpiration) all contributed significantly to determine T/ET (*p <* 0.05; see Fig. 4). T/ET increased with net radiation, precipitation and fAPAR and decreased with air temperature and transpiration. According to the theory of Zhang et al. (2010), higher fAPAR leads to greater energy interception by the canopy and therefore a higher T/ET. More energy (higher *R*_n_ and *T*_a_) and precipitation should, all else equal, lead to a denser canopy and thus larger fAPAR. But increasing temperature increases soil evaporation more than transpiration. The net effect of *T*_a_ on T/ET ratio is therefore negative (Fig. 4d). As T increases, T/ET decreases. This negative effect (Fig. 4e) suggests that soil evaporation is more sensitive to changing energy inputs, due to the limitation of transpiration by stomata.

**Table 3.**
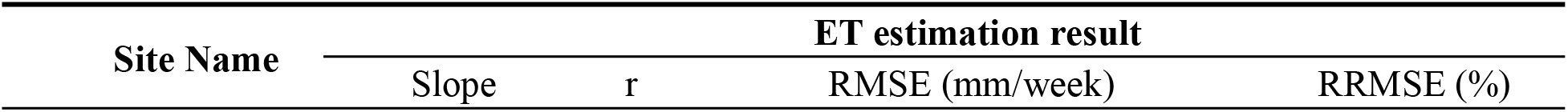
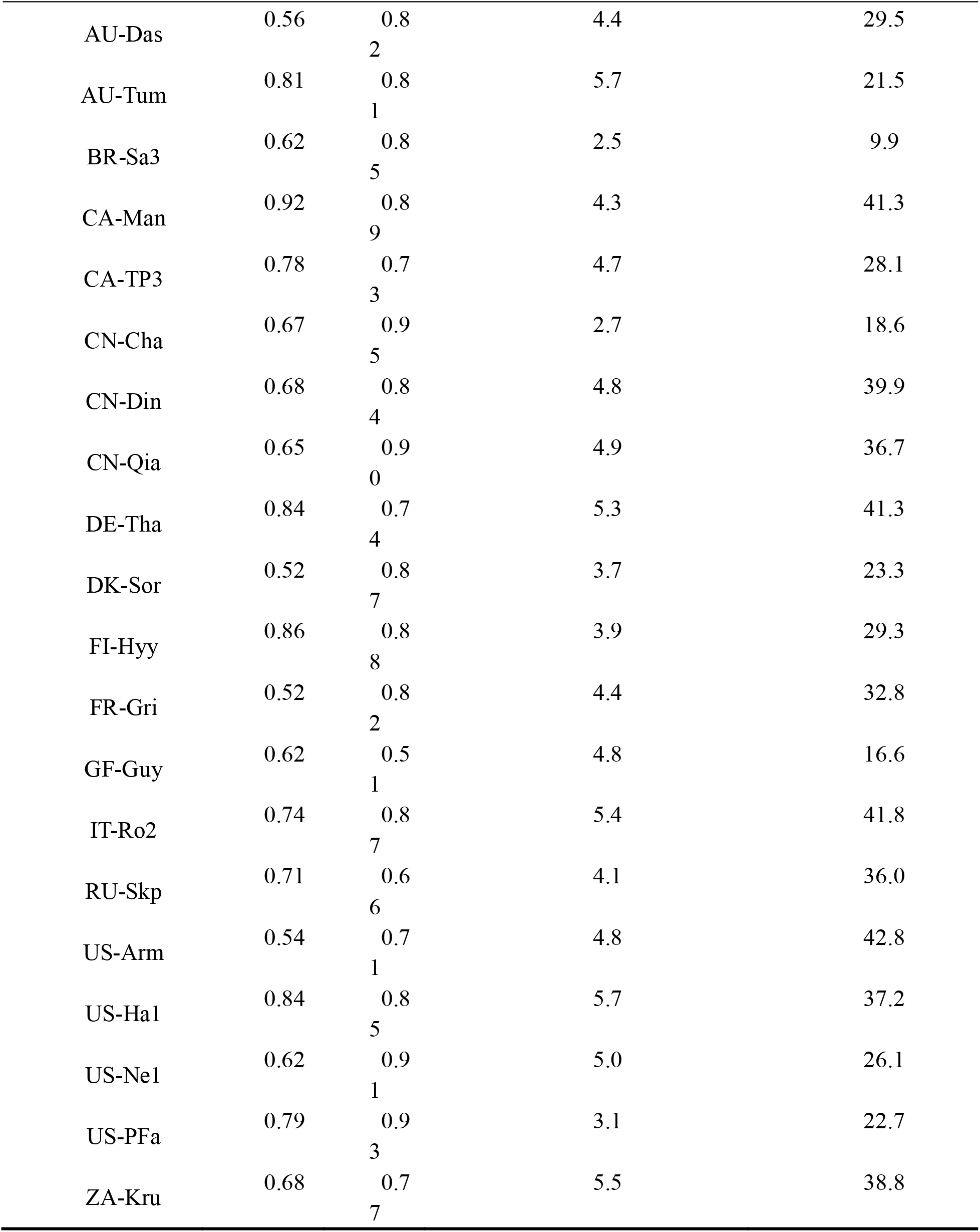
Statistical information on the ET evaluations on a weekly timestep. Slope in table represents linear fitting slope of estimated transpiration to ET observation, which is considered as the T/ET ratio. Other items have the same meaning with its in Table 2.

| Site Name | ET estimation result |  |  |  |
| --- | --- | --- | --- | --- |
|  | Slope | r | RMSE (mm/week) | RRMSE (%) |
| AU-Das | 0.56 | 0.8<br>2 | 4.4 | 29.5 |
| AU-Tum | 0.81 | 0.8<br>1 | 5.7 | 21.5 |
| BR-Sa3 | 0.62 | 0.8<br>5 | 2.5 | 9.9 |
| CA-Man | 0.92 | 0.8<br>9 | 4.3 | 41.3 |
| CA-TP3 | 0.78 | 0.7<br>3 | 4.7 | 28.1 |
| CN-Cha | 0.67 | 0.9<br>5 | 2.7 | 18.6 |
| CN-Din | 0.68 | 0.8<br>4 | 4.8 | 39.9 |
| CN-Qia | 0.65 | 0.9<br>0 | 4.9 | 36.7 |
| DE-Tha | 0.84 | 0.7<br>4 | 5.3 | 41.3 |
| DK-Sor | 0.52 | 0.8<br>7 | 3.7 | 23.3 |
| FI-Hyy | 0.86 | 0.8<br>8 | 3.9 | 29.3 |
| FR-Gri | 0.52 | 0.8<br>2 | 4.4 | 32.8 |
| GF-Guy | 0.62 | 0.5<br>1 | 4.8 | 16.6 |
| IT-Ro2 | 0.74 | 0.8<br>7 | 5.4 | 41.8 |
| RU-Skp | 0.71 | 0.6<br>6 | 4.1 | 36.0 |
| US-Arm | 0.54 | 0.7<br>1 | 4.8 | 42.8 |
| US-Ha1 | 0.84 | 0.8<br>5 | 5.7 | 37.2 |
| US-Ne1 | 0.62 | 0.9<br>1 | 5.0 | 26.1 |
| US-PFa | 0.79 | 0.9<br>3 | 3.1 | 22.7 |
| ZA-Kru | 0.68 | 0.7<br>7 | 5.5 | 38.8 |

**Fig. 4.**
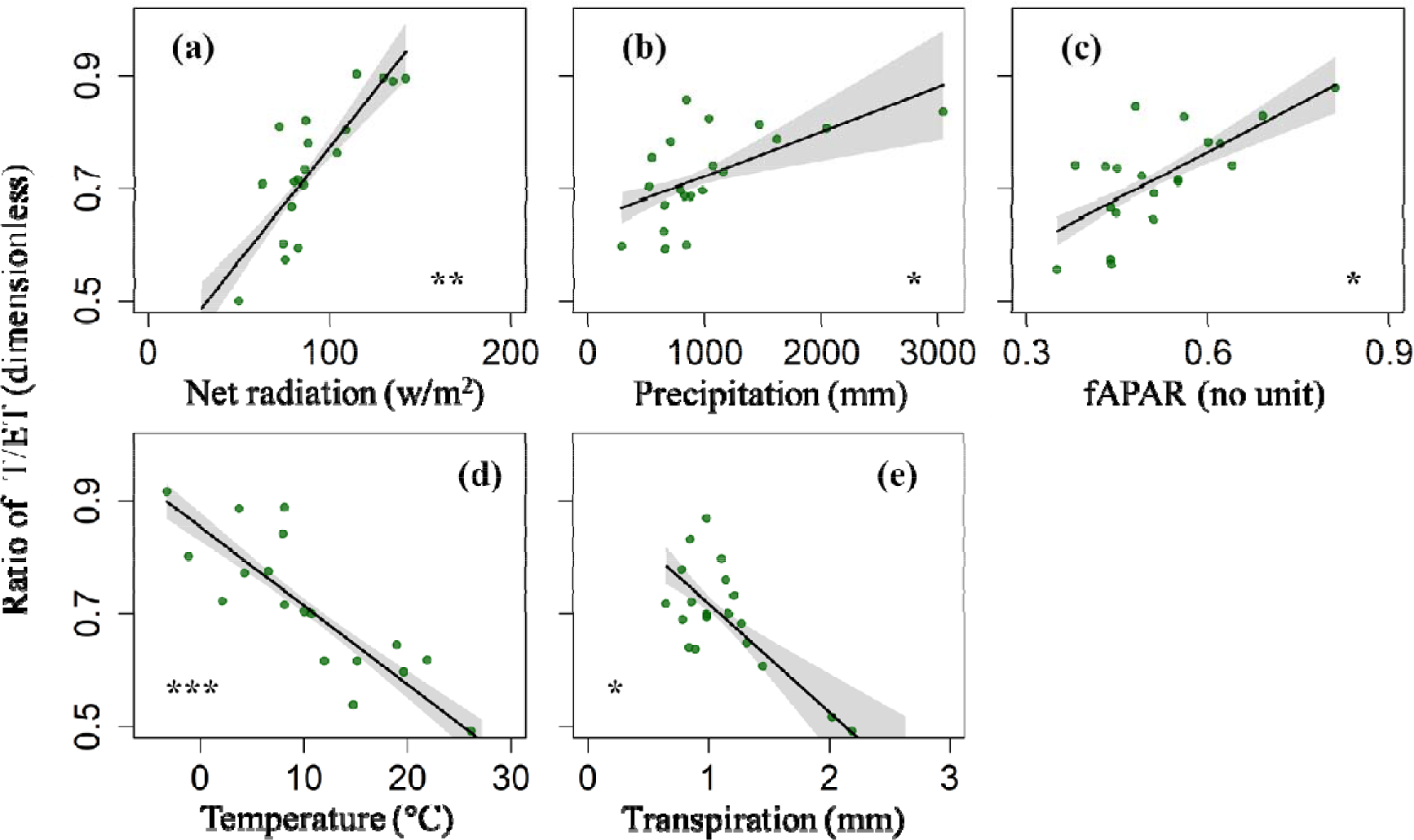
Partial residual plots from the regression of site-scale transpiration to evapotranspiration ratio (T/ET) against environmental predictors. Net radiation, fAPAR, air temperature and modelled transpiration are averaged by effective observation during growing season at each site, except precipitation (mm) in (b) which is an annual accumulated total. Radiation (W/m^2^) in (a) represents net radiation. Grey colours in all panels represents 95% confidence intervals. Significance: *** *p* < 0.001, ** *p* < 0.01, * *p* < 0.05.

We estimated T/ET ratio at site level using the empirical relationship between the T/ET and the five variables using the following relationship:

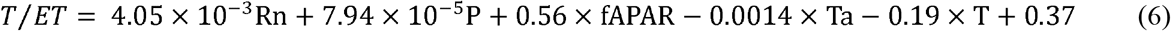

The correlation coefficient of the model against fitted (linear regressed by modelled T against ET observation) T/ET ratio was 0.84 based on site-average conditions (see Fig. 5). Analysis of the dependencies of this estimated ratio on several predictor variables suggested that denser canopies significantly lead to higher values of T/ET, as also suggested by Wang et al. (2014). Results of cross-validation (at each target site re-fitting the relationship using data from other sites, then using the new function to estimate ET at the target site, see Fig. S1) indicate the robustness and reliability of this approach at different sites.

**Fig. 5.**
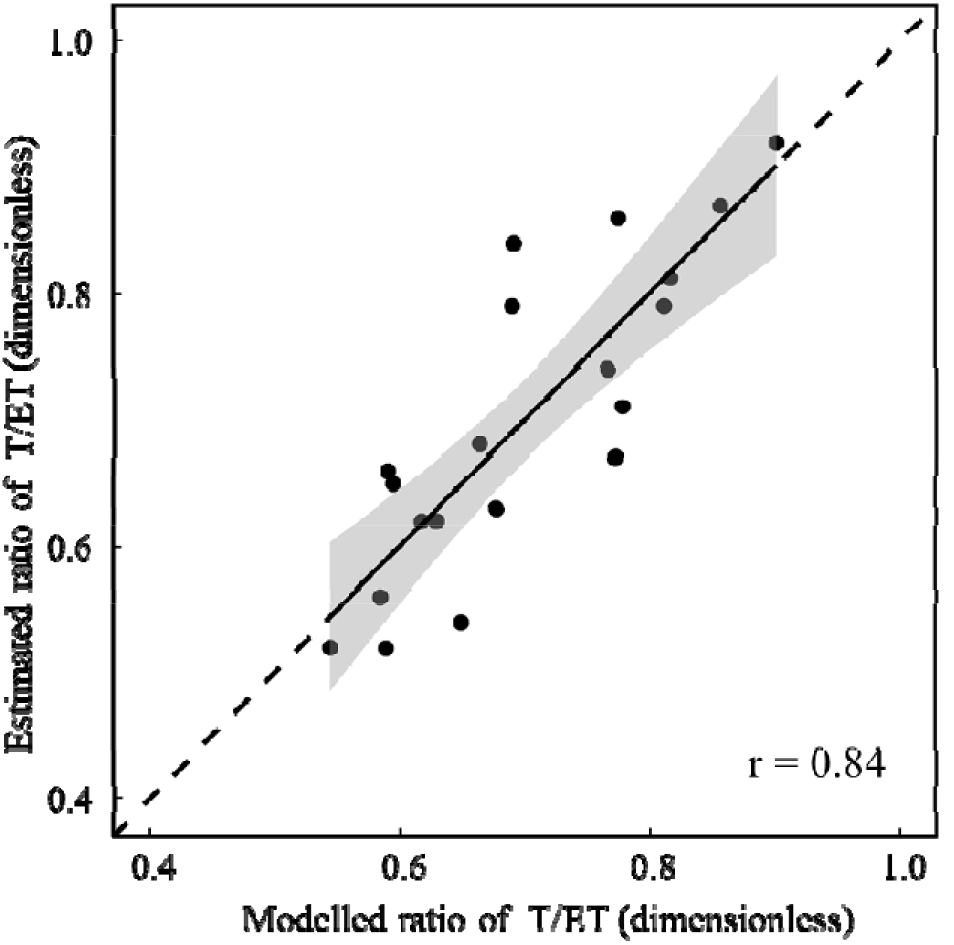
Comparison of predicted and modelled (linear regressed by modelled T against ET observation) T/ET ratio at each flux sites. The predictions are based on the empirical environmental dependences. Grey colour represents 95% confidence intervals; the dashed line is the 1:1 line.

### 3.3 Performance of ET estimation

We obtained a good validation result of modelled ET against flux observations (slope = 0.92, r = 0.85, RMSE = 5.46 mm week^−1^ and NSE = 0.66, see Fig. 6). Good performance of the model was also shown by correlation at site level (*r* ranges from 0.51 to 0.95), and RMSE (from 2.50 to 5.71 mm week^−1^).

**Fig. 6.**
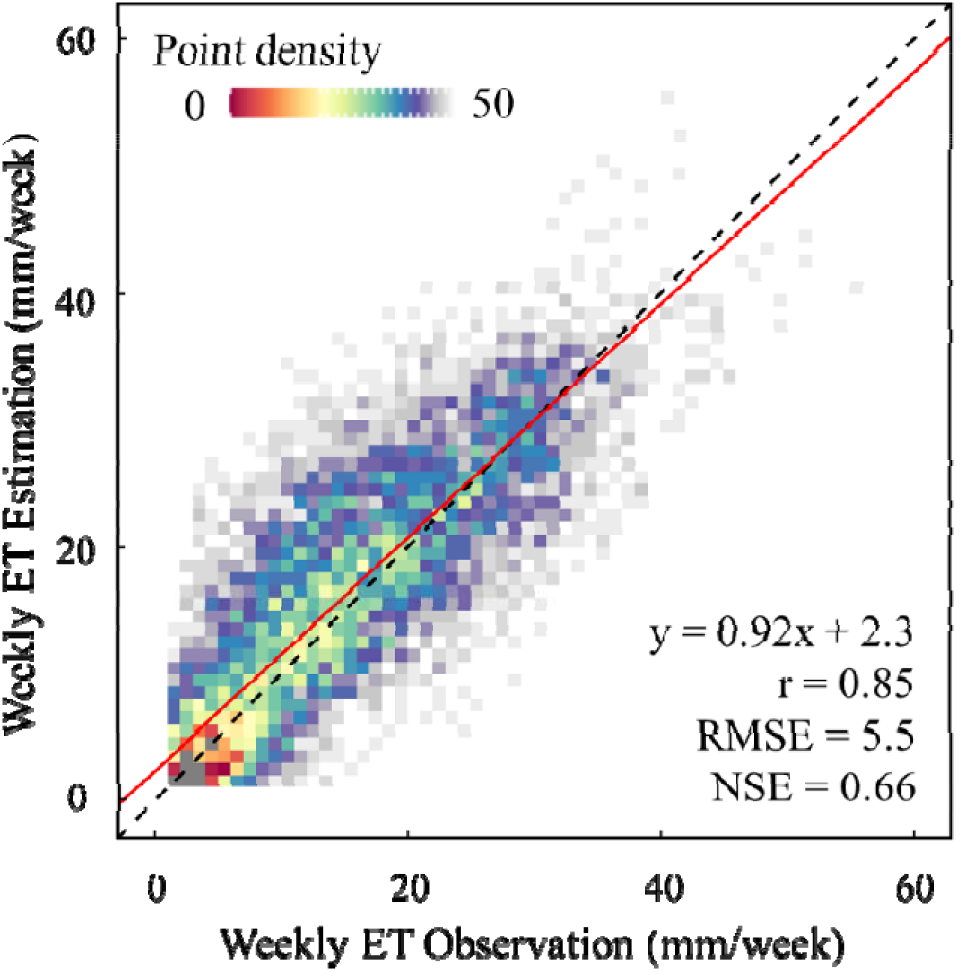
Comparison of estimated ET using the approach in this study against observation with weekly time scale. The red line is the linear fit; the dashed line is the 1:1 line. Colours represents the density of points.

## 4 Discussion

We have demonstrated an approach to estimate evapotranspiration that greatly reduces the need to specify uncertain parameters. It points the way towards a parsimonious general theory of ET estimation from remotely sensed observations and environmental variables. The improvement also avoids discontinuities due to the need to impose biome boundaries when biome-specific parameters are required. This approach, based in part on optimality theory, could provide a means to improve the monitoring and forecasting of ET.

### 4.1 Success of the a priori method

In our framework, GPP and ET are estimated without the need to estimate or calibrate biome or plant functional type specific parameters. The imposition of biome boundaries, required by many existing methods, creates uncertainty for two reasons. First is the quality of terrestrial classification. Remotely sensed classification maps always have reasonable accuracy for types with clear spatial or temporal pattern, but there are still uncertainties, especially in wet regions with major cloud contamination (Zhang et al., 2018). Incorrect biome identification would influence the parameters and thus the results (see the example Fig. S2). In addition, separate parameter fitting by biomes can obscure systematic differences among modelled responses to environment (see Fig. S3). The P model, in contrast, is based on a universal plant optimality theory that implicitly predicts differences among biomes as a consequence of the environments where they occur.

### 4.2 Uncertainties

The P model provides a universal strategy to calculate canopy conductance in C_3_ vegetation and is extended to C_4_ plants in a straightforward way. The good results at the maize site US-Ne1 support this approach. The current version however adopts a constant value of 0.45 for *χ* in C_4_ plants (Farquhar et al., 1989), a simplification that could be improved on.

The conversion from transpiration to total evapotranspiration is based on a T/ET ratio, which was considered as a constant at each site. In Fig. S4, we tested this assumption by constructing a global map of the standard deviation of the annual T/ET ratio in the PMLv2 (Penman◻Monteith-Leuning) data set from 2003 to 2017 (Zhang et al., 2019). PML was chosen because its strategy for partitioning available energy in the PM equation is the same as the one we used. Most of the region (nearly 80%) has a T/ET standard deviation < 0.05 over these 15 years. Less than 1% of the area has a standard deviation > 0.1. In addition, since the energy intercepted by canopy is calculated by fAPAR, area with significant variation of T/ET are mainly located in the region with a significant variation in terrestrial LAI, both greening and browning, as shown by Zhu et al., (2016).

A comparison with global patterns in other global T/ET products, including PML and GLEAM, shows that the three remote-sensing based T/ET estimates yield similar latitudinal profiles (see Fig. S5). However, the averaged value in PML (0.46) is substantially lower than in GLEAM (0.62) and our empirical method (0.68). In regions with extremely sparse vegetation, such as North Africa and the Tibetan plateau (about 0.3 by our method, compared with near-zero in GLEAM and no value given in PML), we overestimated T/ET. There are two possible reasons for this. First, fAPAR has a paramount contribution to our prediction, which responds to the low fAPAR in such regions. Second, we employed only 20 sites during the fitting, and none of them were from arid regions, so there is a possibility of bias.

Comparisons in this study (Fig. S6) and Gu et al., (2018) point to large difference between products and estimation strategies for T/ET. Large differences have also been reported in ESMs (Lian et al., 2018). We hope to improve the performance and temporal resolution of T/ET modelling with the help of multiple observation strategies in near future. Possible more explicitly process-based approaches to modelling the non-T components of ET include the flux variance similarity (FVS) partitioning method proposed by Scanlon and Sahu (2008), and the use of data from lysimeters and sap flow meters (Zhou et al., 2018, Nelson et al., 2020).

The scientific advance presented here lends itself naturally to application with remote sensing technology. The only satellite-derived input is fAPAR, here at 5 km spatial resolution – but this implies a mismatch with the flux tower footprints that may explain the underestimation of GPP at some sites. Although this underestimation is corrected automatically in the process of estimating T/ET ratios, it would be desirable to reduce it by using remotely sensed input with finer spatial resolution, such as images from MODIS (MOderate Resolution Imaging Spectroradiometer), to better fit the tower footprint. There is also the potential to improve the accuracy of fAPAR estimation from satellites. For example, according to Zhang et al. (2020), light absorbed by chlorophyll (Chl) should provide a better basis for the estimation of GPP; a variety of indices that more accurately represent Chl absorption are becoming available. Moreover, ET could usefully be mapped with, say, 30 m or finer spatial resolution, as climate variables tend to vary much more smoothly than vegetation properties (Tan et al., 2019). An improved, high-resolution satellite-based ET estimation strategy would have many applications in land-surface modelling and water resources research.

## Acknowledgements

This research was supported by the National Key R&D Program of China (no. 2018YFA0605400), the National Natural Science Foundation of China (grant no. 91837312, 42001356) and the generosity of Eric and Wendy Schmidt by recommendation of the Schmidt Futures program. This research contributes to the Imperial College initiative on Grand Challenges in Ecosystems and the Environment. ICP’s contribution has been supported by the European Research Council (ERC) under the European Union’s Horizon 2020 research and innovation programme (grant agreement No: 787203 REALM). We used ‘free and fair use’ eddy-covariance data acquired by the FLUXNET community and, in particular, by the following networks: AmeriFlux, ChinaFlux, OzFlux, CarboEuropeIP and TCOS-Siberia. We acknowledge the financial support to the eddy-covariance data harmonization provided by University of Western Australia, Charles Darwin University, CSIRO, University of California – Irvine, University of Manitoba, McMaster University, IGSNRR-, SCIB- and IAE- Chinese Academy of Sciences, Technical University of Denmark, University of Heisinki, INRA Grignon, INRA, University of Tuscia Viterbo, IBPC Russia, Lawrence Berkeley National Laboratory, Harvard University, University of Nebraska- Lincoln, University of Wisconsin and University of the Witwatersrand.

## Supplementary Information

**Fig. S1.**
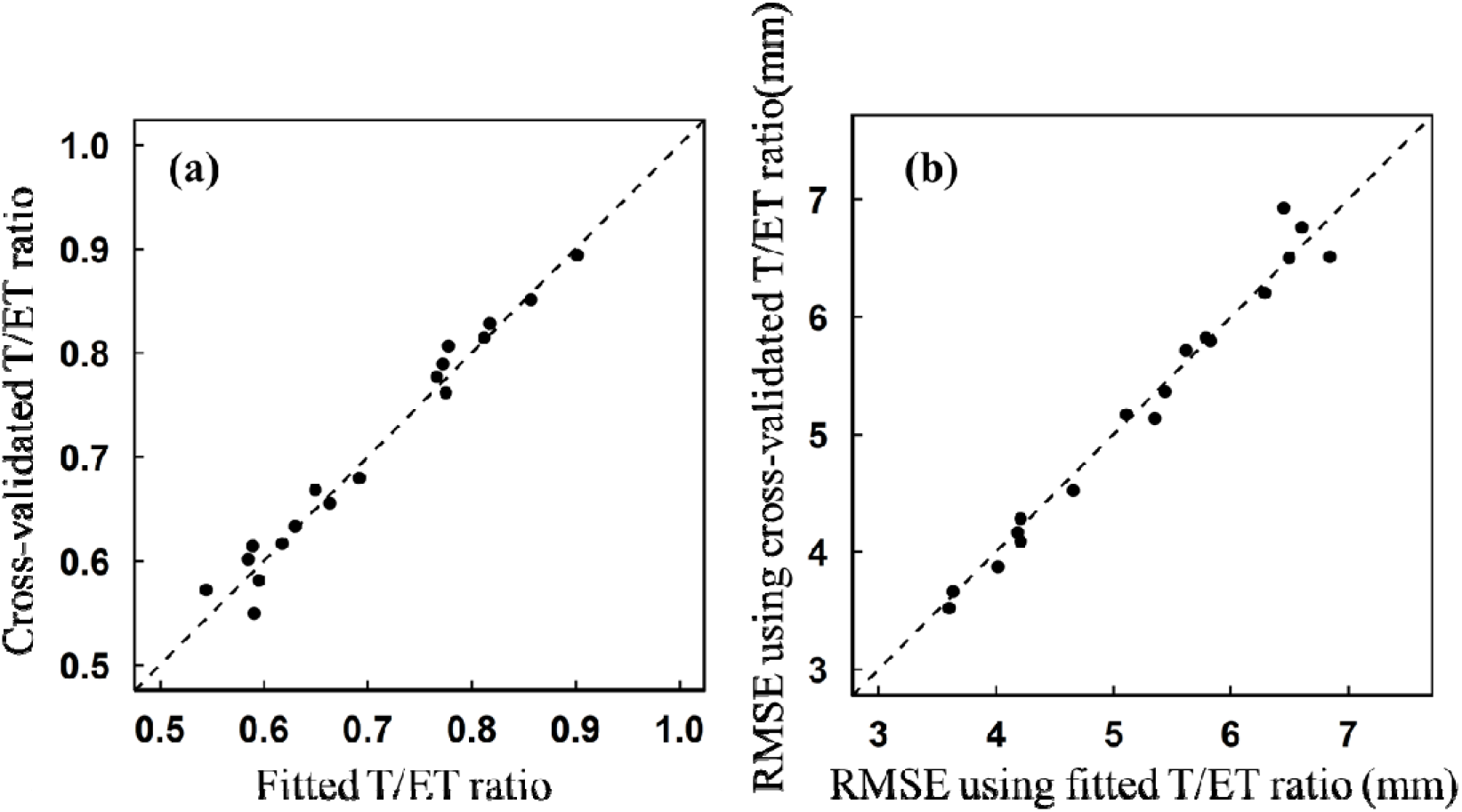
Cross-validation of the T/ET fitting theory. X-axis is fitted T/ET ratio (in panel a) and RMSE of ET result (in panel b) using the universal function. Y-axis is T/ET ratio (in panel a) and RMSE of ET result (in panel b) result using the cross-validated function (exempting each target site during fitting).

**Fig. S2.**
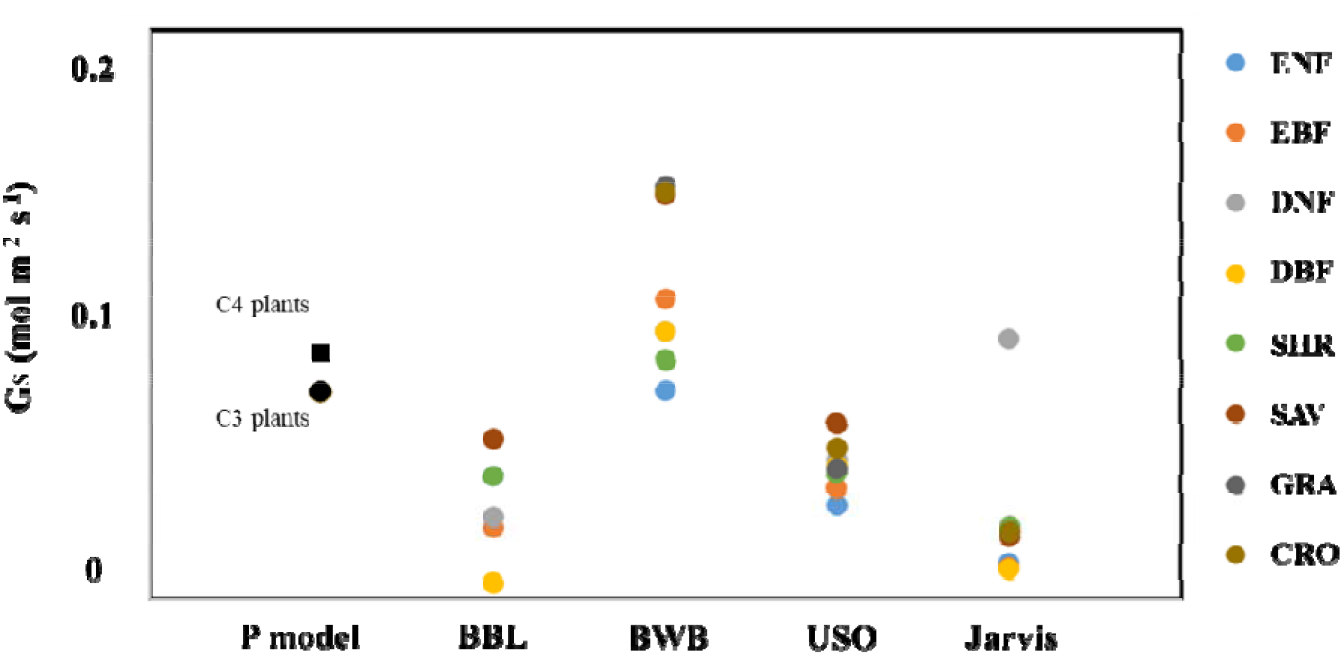
Comparison of typical conductance models under baseline condition. (=20, VPD=10hPa, fAPAR=0.8, =380 μmol mol^−1^, SWC=0.2 m^3^ m^−3^, PPFD=275 μmol^−2^ photon s^−1^). Two black points represents result estimated by C3 and C4 P model version respectively. Different colour represents different biomes types. BBL represents Ball-Berry-Leuning conductance model (Leuning, 1995), BWB represents Ball-Berry model (Ball and Berry, 1987), USO represents Mdelyn model (Medlyn et al., 2011), Jarvis represents Jarvis model (Jarvis, 1976). For each conductance model requires GPP as input, we use the result estimated by P model.

**Fig. S3.**
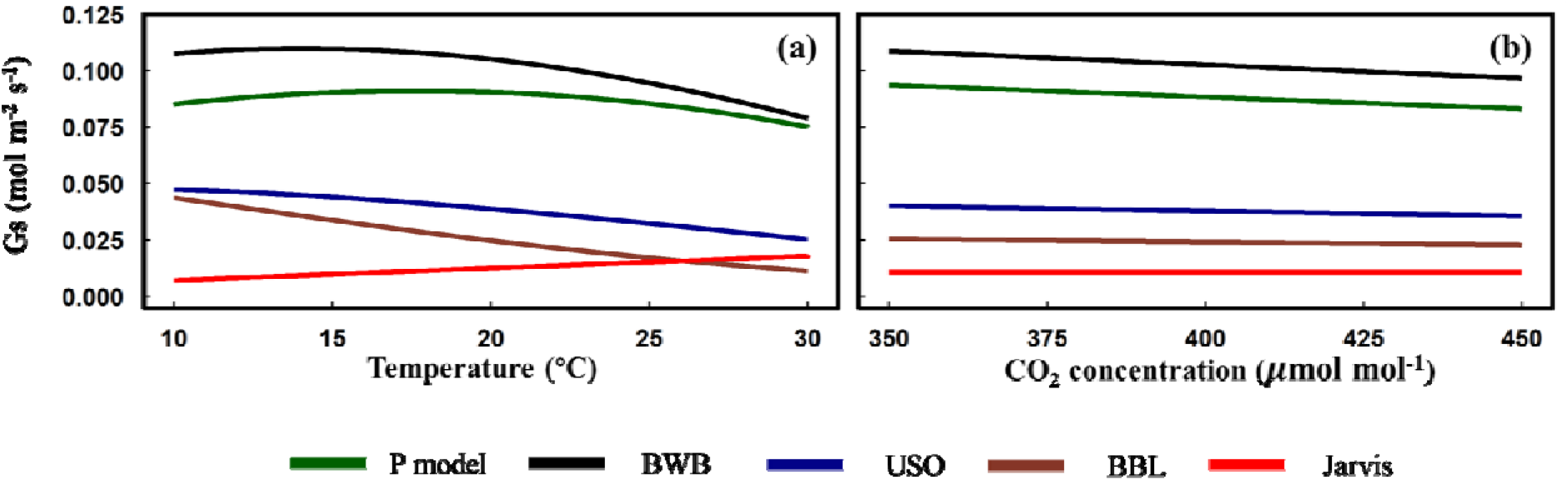
Comparison of typical conductance models response to increasing temperature and ambient CO_2_ concentration. Baseline condition and model code is the same with Fig. S1.

**Fig. S4.**
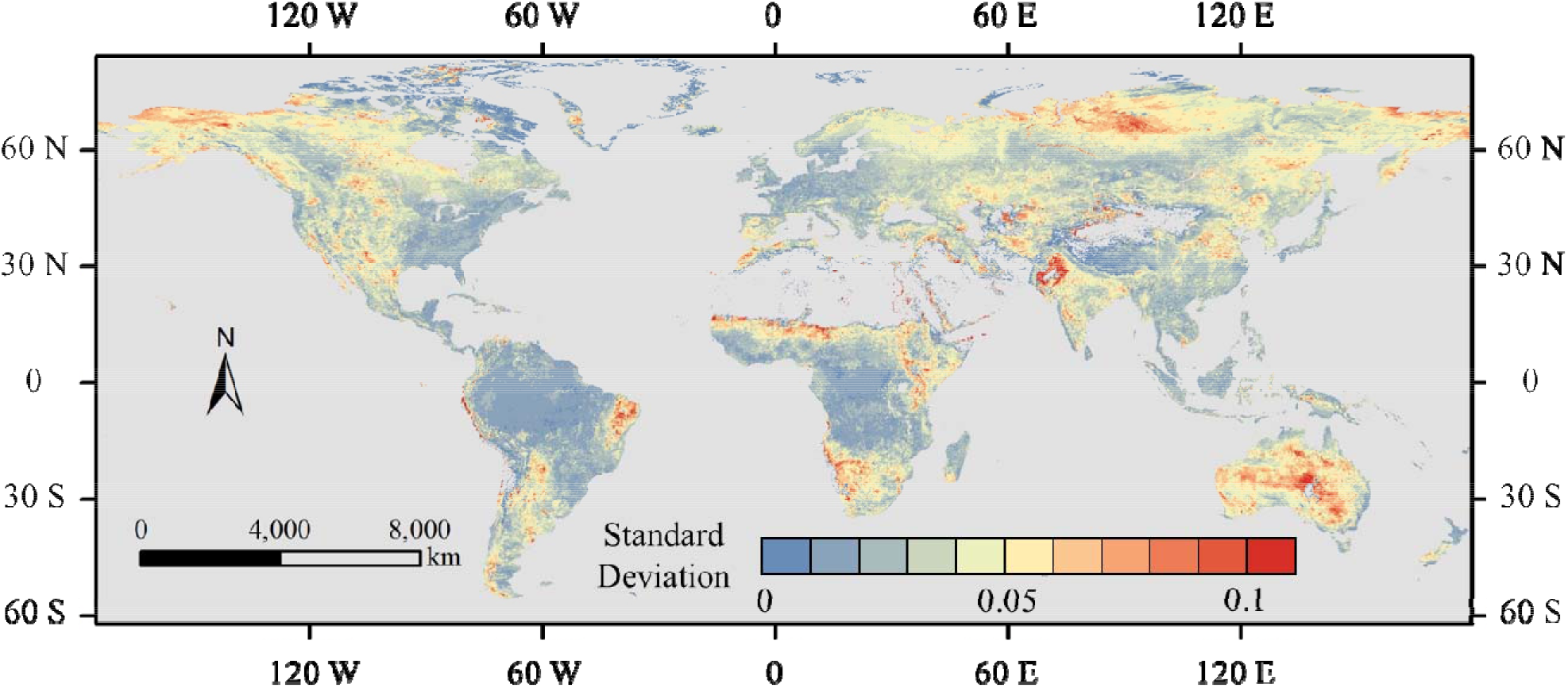
Standard deviation map of PML T/ET ratio. The calculation is based on the PML v2 product from 2003 to 2017 (Zhang et al., 2019).

**Fig. S5.**
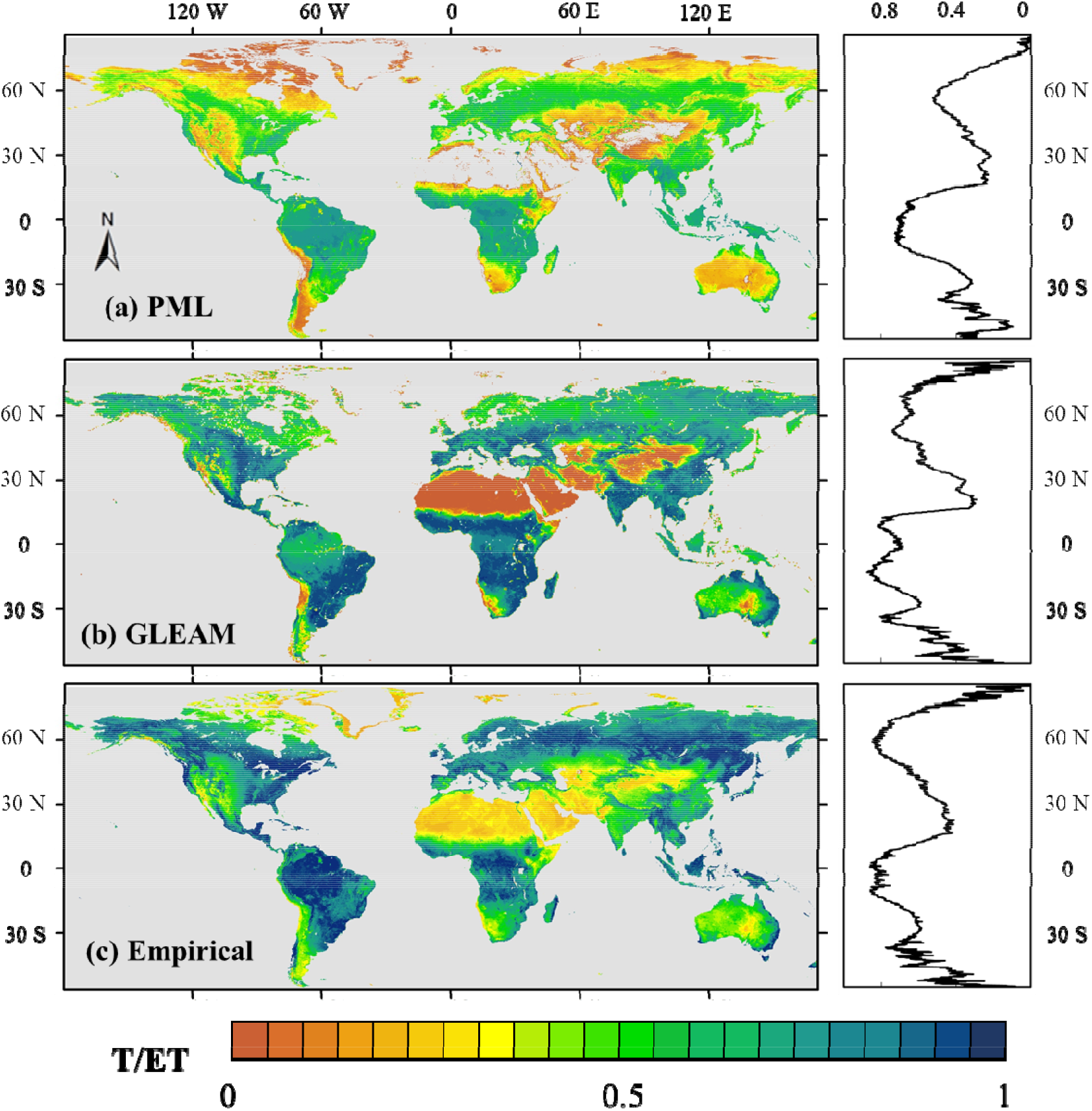
Global distribution of T/ET ratio, provided by (a) PML, (b) GLEAM, and (c) the empirical method in this research. Latitude T/ET profile of each product is given next to each map. Data displayed here is an average between 2003 to 2017, during the period the PML product is available.

**Fig. S6.**
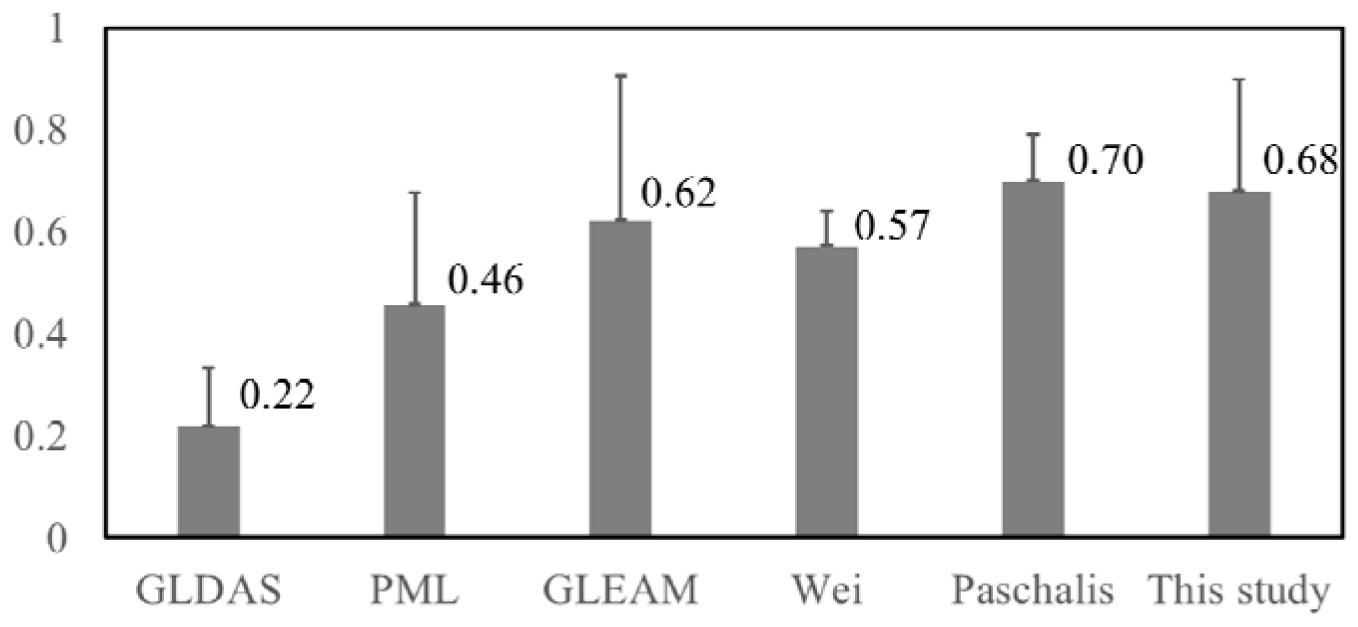
Summarized result of global T/ET ratio. Mean T/ET value is labelled next to each column. Standard deviation is represented by the error bar. Result of Wei was reported by Wei et al., (2017). Result of Paschalis was reported by Paschalis et al., (2018). Global products (GLDAS v2.1, PML v2 and GLEAM v3.3) are calculated based on data between 2003 to 2017.

## Notes

### Competing Interest Statement

The authors have declared no competing interest.

## References

Allen R G, Pereira L S, Raes D, et al. Crop evapotranspiration-Guidelines for computing crop water requirements-FAO Irrigation and drainage paper 56[J]. Fao, Rome, 1998, 300(9): D05109.

Bai P, Liu X. Intercomparison and evaluation of three global high-resolution evapotranspiration products across China[J]. Journal of hydrology, 2018, 566: 743–755.

Ball J T, Woodrow I E, Berry J A. A model predicting stomatal conductance and its contribution to the control of photosynthesis under different environmental conditions[M]//Progress in photosynthesis research. Springer, Dordrecht, 1987: 221–224.

Bernacchi C J, Bagley J E, Serbin S P, et al. Modelling C 3 photosynthesis from the chloroplast to the ecosystem[J]. Plant, Cell & Environment, 2013, 36(9): 1641–1657.

Bastiaanssen W G M, Menenti M, Feddes R A, et al. A remote sensing surface energy balance algorithm for land (SEBAL). 1. Formulation[J]. Journal of hydrology, 1998, 212: 198–212.

Bastiaanssen W G M, Molden D J, Makin I W. Remote sensing for irrigated agriculture: examples from research and possible applications[J]. Agricultural water management, 2000, 46(2): 137–155.

Cai W, Prentice I C. Recent trends in gross primary production and their drivers: analysis and modelling at flux-site and global scales[J]. Environmental Research Letters, 2020.

Carlson T N. Modeling stomatal resistance: an overview of the 1989 workshop at the Pennsylvania State University[J]. Agricultural and Forest Meteorology, 1991, 54(2-4): 103–106.

Chen J L, Reynolds J F, Harley P C, et al. Coordination theory of leaf nitrogen distribution in a canopy[J]. Oecologia, 1993, 93(1): 63–69.

Chen J, Jönsson P, Tamura M, et al. A simple method for reconstructing a high-quality NDVI time-series data set based on the Savitzky–Golay filter[J]. Remote sensing of Environment, 2004, 91(3-4): 332–344.

Cheng L, Zhang L, Wang Y P, et al. Recent increases in terrestrial carbon uptake at little cost to the water cycle[J]. Nature Communications, 2017, 8(1): 1–10.

Claverie M, Vermote E. NOAA Climate Data Record (CDR) of Leaf Area Index (LAI) and Fraction of Absorbed Photosynthetically Active Radiation (FAPAR) Version 4[J]. NOAA National Centers for Environmental Information, 2014.

Cleugh H A, Leuning R, Mu Q, et al. Regional evaporation estimates from flux tower and MODIS satellite data[J]. Remote Sensing of Environment, 2007, 106(3): 285–304.

Collatz G J, Ribas-Carbo M, Berry J A. Coupled photosynthesis-stomatal conductance model for leaves of C4 plants[J]. Functional Plant Biology, 1992, 19(5): 519–538.

Cowan I R, Farquhar G D, Stomatal function in relation to leaf metabolism and environment[J]. 1977.

Elnashar A, Wang L, Wu B, et al. Synthesis of global actual evapotranspiration from 1982 to 2019[J]. Earth System Science Data, 2021, 13(2): 447–480.

Ershadi A, McCabe M F, Evans J P, et al. Multi-site evaluation of terrestrial evaporation models using FLUXNET data[J]. Agricultural and Forest Meteorology, 2014, 187: 46–61.

Farquhar G D, von Caemmerer S, Berry J A. A biochemical model of photosynthetic CO 2 assimilation in leaves of C 3 species[J]. Planta, 1980, 149(1): 78–90.

Farquhar G D, Ehleringer J R, Hubick K T. Carbon isotope discrimination and photosynthesis[J]. Annual review of plant biology, 1989, 40(1): 503–537.

Fatichi S, Pappas C. Constrained variability of modeled T: ET ratio across biomes[J]. Geophysical Research Letters, 2017, 44(13): 6795–6803.

Fisher J B, Melton F, Middleton E, et al. The future of evapotranspiration: Global requirements for ecosystem functioning, carbon and climate feedbacks, agricultural management, and water resources[J]. Water Resources Research, 2017, 53(4): 2618–2626.

Franklin O, Harrison S P, Dewar R, et al. Organizing principles for vegetation dynamics[J]. Nature plants, 2020: 1–10.

Gallego-Sala A V, Charman D J, Brewer S, et al. Latitudinal limits to the predicted increase of the peatland carbon sink with warming[J]. Nature climate change, 2018, 8(10): 907–913.

Gan R, Zhang Y, Shi H, et al. Use of satellite leaf area index estimating evapotranspiration and gross assimilation for Australian ecosystems[J]. Ecohydrology, 2018, 11(5): e1974.

Gitelson A A, Arkebauer T J, Suyker A E. Convergence of daily light use efficiency in irrigated and rainfed C3 and C4 crops[J]. Remote sensing of environment, 2018, 217: 30–37.

Granger R J, Gray D M. Evaporation from natural non saturated surfaces[J]. J. Hydrol, 1989, 11121: 29.

Gu C, Ma J, Zhu G, et al. Partitioning evapotranspiration using an optimized satellite-based ET model across biomes[J]. Agricultural and forest meteorology, 2018, 259: 355–363.

Haxeltine, A. & Prentice, I.C. (1996). A general model for the light-use efficiency of primary production. Functional Ecology, 10, 551–561.

He L, Chen J M, Gonsamo A, et al. Changes in the shadow: the shifting role of shaded leaves in global carbon and water cycles under climate change[J]. Geophysical Research Letters, 2018, 45(10): 5052–5061.

Jarvis P G. The interpretation of the variations in leaf water potential and stomatal conductance found in canopies in the field[J]. Philosophical Transactions of the Royal Society of London. B, Biological Sciences, 1976, 273(927): 593–610.

Jiang C, Ryu Y. Multi-scale evaluation of global gross primary productivity and evapotranspiration products derived from Breathing Earth System Simulator (BESS)[J]. Remote Sensing of Environment, 2016, 186: 528–547.

Jung M, Reichstein M, Bondeau A. Towards global empirical upscaling of FLUXNET eddy covariance observations: validation of a model tree ensemble approach using a biosphere model[J]. 2009.

Kljun N, Calanca P, Rotach M W, et al. A simple two-dimensional parameterisation for Flux Footprint Prediction (FFP)[J]. Geoscientific Model Development, 2015, 8(11): 3695.

Knauer J, Zaehle S, Medlyn B E, et al. Towards physiologically meaningful water◻use efficiency estimates from eddy covariance data[J]. Global Change Biology, 2018, 24(2): 694–710.

Kool D, Agam N, Lazarovitch N, et al. A review of approaches for evapotranspiration partitioning[J]. Agricultural and forest meteorology, 2014, 184: 56–70.

Kowalczyk E A, Wang Y P, Law R M, et al. The CSIRO Atmosphere Biosphere Land Exchange (CABLE) model for use in climate models and as an offline model[J]. CSIRO Marine and Atmospheric Research Paper, 2006, 13: 42.

Kubien D S, von Caemmerer S, Furbank R T, et al. C4 photosynthesis at low temperature. A study using transgenic plants with reduced amounts of Rubisco[J]. Plant Physiology, 2003, 132(3): 1577–1585.

Lagouarde J P. Use of NOAA AVHRR data combined with an agrometeorological model for evaporation mapping[J]. International Journal of Remote Sensing, 1991, 12(9): 1853–1864.

Leuning R. A critical appraisal of a combined stomatal◻photosynthesis model for C3 plants[J]. Plant, Cell & Environment, 1995, 18(4): 339–355.

Leuning R, Zhang Y Q, Rajaud A, et al. A simple surface conductance model to estimate regional evaporation using MODIS leaf area index and the Penman◻Monteith equation[J]. Water Resources Research, 2008, 44(10).

Lian X, Piao S, Huntingford C, et al. Partitioning global land evapotranspiration using CMIP5 models constrained by observations[J]. Nature Climate Change, 2018, 8(7): 640–646.

Lin Y S, Medlyn B E, Duursma R A, et al. Optimal stomatal behaviour around the world[J]. Nature Climate Change, 2015, 5(5): 459–464.

Maire V, Martre P, Kattge J, et al. The coordination of leaf photosynthesis links C and N fluxes in C 3 plant species[J]. PloS one, 2012, 7(6): e38345.

Medlyn B E, Duursma R A, Eamus D, et al. Reconciling the optimal and empirical approaches to modelling stomatal conductance[J]. Global Change Biology, 2011, 17(6): 2134–2144.

Moran M S, Clarke T R, Inoue Y, et al. Estimating crop water deficit using the relation between surface-air temperature and spectral vegetation index[J]. Remote sensing of environment, 1994, 49(3): 246–263.

Mu Q, Heinsch F A, Zhao M, et al. Development of a global evapotranspiration algorithm based on MODIS and global meteorology data[J]. Remote sensing of Environment, 2007, 111(4): 519–536.

Nagler P L, Glenn E P, Nguyen U, et al. Estimating riparian and agricultural actual evapotranspiration by reference evapotranspiration and MODIS enhanced vegetation index[J]. Remote Sensing, 2013, 5(8): 3849–3871.

Nelson J A, Pérez◻Priego O, Zhou S, et al. Ecosystem transpiration and evaporation: Insights from three water flux partitioning methods across FLUXNET sites[J]. Global change biology, 2020, 26(12): 6916–6930.

Norman J M, Kustas W P, Humes K S. Source approach for estimating soil and vegetation energy fluxes in observations of directional radiometric surface temperature[J]. Agricultural and Forest Meteorology, 1995, 77(3-4): 263–293.

Oleson K W, Lawrence D M, Gordon B, et al. Technical description of version 4.0 of the Community Land Model (CLM)[J]. 2010.

Paschalis A, Fatichi S, Pappas C, et al. Covariation of vegetation and climate constrains present and future T/ET variability[J]. Environmental Research Letters, 2018, 13(10): 104012.

Perez◻Priego O, Katul G, Reichstein M, et al. Partitioning eddy covariance water flux components using physiological and micrometeorological approaches[J]. Journal of Geophysical Research: Biogeosciences, 2018, 123(10): 3353–3370.

Prentice I C, Dong N, Gleason S M, et al. Balancing the costs of carbon gain and water transport: testing a new theoretical framework for plant functional ecology[J]. Ecology letters, 2014, 17(1): 82–91.

Priestley C H B, Taylor R J. On the assessment of surface heat flux and evaporation using large-scale parameters[J]. Monthly weather review, 1972, 100(2): 81–92.

Qiao S, Wang H, Prentice I C, et al. Extending a first-principles primary production model to predict wheat yields[J]. Agricultural and Forest Meteorology, 2020, 287: 107932.

Reichstein M, Falge E, Baldocchi D, et al. On the separation of net ecosystem exchange into assimilation and ecosystem respiration: review and improved algorithm[J]. Global change biology, 2005, 11(9): 1424–1439.

Rodell M, Houser P R, Jambor U E A, et al. The global land data assimilation system[J]. Bulletin of the American Meteorological Society, 2004, 85(3): 381–394.

Sage R F. The evolution of C4 photosynthesis[J]. New phytologist, 2004, 161(2): 341–370.

Sayre R T, Kennedy R A, Pringnitz D J. Photosynthetic enzyme activities and localization in Mollugo verticillata populations differing in the levels of C3 and C4 cycle operation[J]. Plant Physiology, 1979, 64(2): 293–299.

Scanlon T M, Sahu P. On the correlation structure of water vapor and carbon dioxide in the atmospheric surface layer: A basis for flux partitioning[J]. Water Resources Research, 2008, 44(10).

Smith N G, Keenan T F, Colin Prentice I, et al. Global photosynthetic capacity is optimized to the environment[J]. Ecology letters, 2019, 22(3): 506–517.

Stocker B D, Zscheischler J, Keenan T F, et al. Quantifying soil moisture impacts on light use efficiency across biomes[J]. New Phytologist, 2018, 218(4): 1430–1449.

Stocker B D, Wang H, Smith N G, et al. P-model v1. 0: an optimality-based light use efficiency model for simulating ecosystem gross primary production[J]. Geoscientific Model Development, 2020, 13(3): 1545–1581.

Su Z. The Surface Energy Balance System (SEBS) for estimation of turbulent heat fluxes[J]. Hydrology and earth system sciences, 2002, 6(1): 85–99.

Tan S, Wu B, Yan N. A method for downscaling daily evapotranspiration based on 30-m surface resistance[J]. Journal of Hydrology, 2019, 577: 123882.

Thom A S. Momentum, mass and heat exchange of vegetation[J]. Quarterly Journal of the Royal Meteorological Society, 1972, 98(415): 124–134.

Torres A F, Walker W R, McKee M. Forecasting daily potential evapotranspiration using machine learning and limited climatic data[J]. Agricultural Water Management, 2011, 98(4): 553–562.

Trenberth K E, Fasullo J T, Kiehl J. Earth’s global energy budget[J]. Bulletin of the American Meteorological Society, 2009, 90(3): 311–324.

Wang H, Prentice I C, Keenan T F, et al. Towards a universal model for carbon dioxide uptake by plants[J]. Nature Plants, 2017, 3(9): 734–741.

Wong S C, Cowan I R, Farquhar G D. Stomatal conductance correlates with photosynthetic capacity[J]. Nature, 1979, 282(5737): 424–426.

Wright, I.J., Reich, P.B. & Westoby, M. (2003). Least◻cost input mixtures of water and nitrogen for photosynthesis. The American Naturalist, 161, 98–111.

Wang L, Good S P, Caylor K K. Global synthesis of vegetation control on evapotranspiration partitioning[J]. Geophysical Research Letters, 2014, 41(19): 6753–6757.

Wei Z, Yoshimura K, Wang L, et al. Revisiting the contribution of transpiration to global terrestrial evapotranspiration[J]. Geophysical Research Letters, 2017, 44(6): 2792–2801.

Wu B, Yan N, Xiong J, et al. Validation of ETWatch using field measurements at diverse landscapes: A case study in Hai Basin of China[J]. Journal of Hydrology, 2012, 436: 67–80.

Yang Y, Roderick M L, Zhang S, et al. Hydrologic implications of vegetation response to elevated CO 2 in climate projections[J]. Nature Climate Change, 2019, 9(1): 44–48.

Zeng H, Wu B, Zhu W, et al. A trade-off method between environment restoration and human water consumption: A case study in Ebinur Lake[J]. Journal of cleaner production, 2019, 217: 732–741.

Zhang X, Wu B, Ponce-Campos G E, et al. Mapping up-to-date paddy rice extent at 10 m resolution in China through the integration of optical and synthetic aperture radar images[J]. Remote Sensing, 2018, 10(8): 1200.

Zhang Y, Leuning R, Hutley L B, et al. Using long◻term water balances to parameterize surface conductances and calculate evaporation at 0.05 spatial resolution[J]. Water Resources Research, 2010, 46(5).

Zhang Y, Kong D, Gan R, et al. Coupled estimation of 500 m and 8-day resolution global evapotranspiration and gross primary production in 2002–2017[J]. Remote sensing of environment, 2019, 222: 165–182.

Zhang Z, Zhang Y, Zhang Y, et al. The potential of satellite FPAR product for GPP estimation: An indirect evaluation using solar-induced chlorophyll fluorescence[J]. Remote Sensing of Environment, 2020, 240: 111686.

Zhou S, Yu B, Zhang Y, et al. Water use efficiency and evapotranspiration partitioning for three typical ecosystems in the Heihe River Basin, northwestern China[J]. Agricultural and Forest Meteorology, 2018, 253: 261–273.

Zhu Z, Piao S, Myneni R B, et al. Greening of the Earth and its drivers[J]. Nature climate change, 2016, 6(8): 791–795.

